# Reactivation of latent infections with migration shapes population-level disease dynamics

**DOI:** 10.1101/2020.05.12.091736

**Authors:** Daniel J. Becker, Ellen D. Ketterson, Richard J. Hall

**Affiliations:** Department of Biology, Indiana University, Bloomington, IN; Center for the Ecology of Infectious Disease, University of Georgia, Athens, GA; Environmental Resilience Institute, Indiana University, Bloomington, IN; Odum School of Ecology, University of Georgia, Athens, GA; Department of Infectious Diseases, College of Veterinary Medicine, University of Georgia, Athens, GA

**Keywords:** animal migration, mechanistic models, recrudescence, migratory relapse, infectious disease

## Abstract

Annual migration is common across animal taxa and can dramatically shape the spatial and temporal patterns of infectious disease. Although migration can decrease infection prevalence in some contexts, these energetically costly long-distance movements can also have immunosuppressive effects that may interact with transmission processes in complex ways. Here we develop a mechanistic model for the reactivation of latent infections driven by physiological changes or energetic costs associated with migration (i.e., “migratory relapse”) and its effects on disease dynamics. We determine conditions under which migratory relapse can amplify or reduce infection prevalence across pathogen and host traits (e.g., infectious periods, virulence, overwinter survival, timing of relapse) and transmission phenologies. We show that relapse at either the start or end of migration can dramatically increase prevalence across the annual cycle and may be crucial for maintaining pathogens with low transmissibility and short infectious periods in migratory populations. Conversely, relapse at the start of migration can reduce the prevalence of highly virulent pathogens by amplifying culling of infected hosts during costly migration, especially for highly transmissible pathogens and those transmitted during migration or the breeding season. Our study provides a mechanistic foundation for understanding the spatiotemporal patterns of relapsing infections in migratory hosts, with implications for zoonotic surveillance and understanding how infection patterns will respond to shifts in migratory propensity associated with environmental change. Further, our work suggests incorporating within-host processes into population-level models of pathogen transmission may be crucial for reconciling the range of migration–infection relationships observed across migratory species.

## Introduction

Long-distance migration is increasingly recognized to shape the spatial and temporal patterns of infectious disease [1,2]. As these seasonal movements between breeding and wintering grounds occur across animals [3], migration can facilitate the geographic spread of zoonotic pathogens such as filoviruses and West Nile virus [4,5]. Pathogens can also threaten migratory hosts, as observed with sockeye salmon and some waterfowl [6,7]. Accordingly, quantifying the conditions under which migration enhances or dampens pathogen transmission is important to identify when and where infection risks in such species are greatest and to predict the epidemiological consequences of shifting migrations with changing land-use and climate [8,9].

Empirical studies have suggested several ecological mechanisms by which animal migration could increase or decrease infection risks. Seasonal movement from breeding to wintering grounds could expose hosts to a greater number of infected individuals, arthropod vectors, or environmental infectious stages across habitats (i.e., migratory exposure [10–12]). However, because long-distance movement is energetically demanding, migration could also reduce infection prevalence by causing high mortality of infected hosts (i.e., migratory culling [13]). Migration could also allow hosts to escape habitats with high infection risk (i.e., migratory escape [14]) or temporally separate from groups of particularly infectious conspecifics (i.e., migratory allopatry [15]). Additionally, certain habitats experienced during migration could also directly cull pathogens sensitive to environmental conditions (i.e., migratory recovery [16]).

Despite theoretical support [17,18] and grounding of transmission-reducing mechanisms in some empirical systems [13,15,19], recent data syntheses suggest animal migration does not uniformly reduce wildlife disease risks. For example, a meta-analysis across host taxa found relatively weak negative effects of infection status and intensity on survival during migration [20], and a comparative study across migrant, resident, and nomadic ungulates also demonstrated little evidence for migratory escape or culling [21]. Similar analyses have also suggested mixed support for migratory exposure to explain greater infection risks with long-distance movement. Specifically, although habitat diversity predicts parasite richness across ungulates, migratory or nomadic species do not sample more diverse habitats than residents [21]. Further, whereas bird species with greater habitat diversity likewise have higher helminth richness, migratory and resident hosts do not systematically differ in habitat use [22]. Such work suggests greater pathogen exposure of migrants may be insufficient to explain observed infection patterns and that other factors related to host migration more likely drive increased infection risks.

One underexplored mechanism focuses on how energetic costs of migration could drive within-host infection processes [23]. Long-distance movement requires substantial energy [24], with some songbirds and temperate bats investing 25–50% of their body mass in fat reserves for flight [25]. Such energy demands and physiological tradeoffs can negatively affect migrant immunity [26–28]. Impaired immunity prior to and during migration could make hosts more susceptible to new infections [29]. However, greater susceptibility would likely increase pathogen transmission only when exposure occurs at these stages of the annual cycle (e.g., breeding grounds before fall migration, wintering grounds in early spring, and stopover sites [11,30]). As an alternative mechanism, immunosuppression associated with migration could cause latent infections (e.g., obtained in a previous season) to reactivate. In a rare experimental test, modified photoperiod of redwings caused the relapse of latent *Borrelia burgdorferi* infections upon initiation of migratory restlessness [31]. Pathogen reactivation caused by physiological preparations for long-distance movement, which we denote “migratory relapse”, could thus facilitate migratory hosts arriving at their breeding grounds and their wintering grounds primed to infect susceptible conspecifics, arthropod vectors, and spillover hosts.

Pathogens that exhibit cycles of latency and reactivation (i.e., susceptible–infected– latent–infected dynamics) are relatively common, particularly for viruses, bacteria, and some protozoa [32–34], and infect migratory hosts ranging from birds to ungulates to bats (Table 1). Immunosuppression from stressors broadly drives the reactivation of pathogens such as herpesviruses in ungulates [35,36], haemosporidians and flaviviruses in songbirds [37,38], and henipaviruses in flying foxes [39,40]. Similarly, migration has been implicated in the relapse of not only *Borrelia burgdorferi* in thrushes but also West Nile virus in white storks [41] and haemosporidians in rusty blackbirds [42], among other examples (Table 1). Recent comparative analyses also found that temperate bird species with a broader range of migratory movements from their wintering grounds (where vector exposure is unlikely) to their breeding grounds are more capable of infecting susceptible ticks, which could be driven by migratory relapse [43]. Such findings more generally underscore that animal migration could play an underrecognized role in the spatial and temporal dynamics of relapsing pathogens across diverse mobile species.

**Table 1.** Examples of pathogens that exhibit cycles of latency and reactivation, their detection in migratory or nomadic host species, and likely drivers of observed relapse events.

| Pathogen | Host taxon | Behavior | Likely driver of relapse | Reference |
| --- | --- | --- | --- | --- |
| *Borrelia burgdorferi* | Redwing<br>(*Turdus iliacus*) | Migrant | Migration | [31] |
| Eastern equine encephalitis virus | Passeriformes | Migrant | Migration, reproduction | [71] |
| *Plasmodium* spp. | Passeriformes | Migrant | Migration, reproduction | [32] |
| West Nile virus | White storks<br>(*Ciconia ciconia*) | Migrant | Migration | [41] |
| Haemosporida | Rusty blackbird<br>(*Euphagus carolinus*) | Migrant | Migration | [42] |
| Infectious hematopoietic necrosis virus | Sockeye salmon<br>(*Oncorhynchus nerka*) | Migrant | Migration | [51] |
| *Brucella abortus* | Elk<br>(*Cervus canadensis*) | Migrant | Reproduction | [61] |
| Equine herpesviruses | Grévy's zebras<br>(*Equus grevyi*) | Migrant | Translocation | [36] |
| Henipaviruses | Flying foxes<br>(*Pteropus* spp.) | Nomad | Reproduction, nutritional stress | [39] |

Mechanistic models of relapsing infections in closed populations broadly suggest that reactivation is an important determinant of pathogen persistence [40], especially when transmission is seasonal (e.g., many vector-borne diseases [44,45]). For migratory species, although pathogen transmission often occurs at the breeding grounds [17], exposure opportunities can take place at other or multiple stages of the annual cycle [11,30,46,47]. If transmission occurs in only the breeding or wintering grounds, relapse during spring and fall migration could facilitate pathogen persistence, akin to biannual birth pulses [48], and elevate prevalence through increasing the force of infection. However, migratory relapse may have strong impacts only when enough hosts are latently infected, suggesting that pathogen traits could moderate when and where relapse increases prevalence. Short infectious periods relative to migration duration could limit how strongly migrants contribute to pathogen transmission upon arrival at the breeding or wintering grounds. Similarly, relapse of particularly virulent pathogens could rapidly cull infected hosts and reduce prevalence [1]. Integrating relapse into general theory for infection dynamics in a migratory host population could accordingly help to disentangle when animal migration is most likely to magnify infectious disease risks.

We here develop a mechanistic model of relapsing infections in a migratory host to explore the conditions under which migration-induced pathogen reactivation amplifies or dampens infection risks. We expand prior modeling frameworks for a host undergoing a two-way annual migration between the breeding and wintering grounds [17,18]. By assuming the start of spring and fall migration causes some fraction of latently infected hosts to relapse, we explore how the frequency of reactivation affects infection dynamics across the annual cycle of a migratory species. Additionally, we identify pathogen traits for which migratory relapse most influences disease risk and assess how the phenology of pathogen transmission modifies infection outcomes. Using models to explore these scenarios could help establish the kinds of host–pathogen systems for which reactivation driven by animal migration can amplify or reduce infection prevalence and thus better guide future empirical tests of model predictions [49].

## Methods

### Model structure

We developed a differential equation model describing population and infection dynamics during breeding, migration, and overwintering periods. We coarsely parameterized our model around a widespread migratory songbird, but our model structure remains general enough to apply to a range of migratory host and pathogen systems in which latent infections reactivate.

#### Host demography and migration

In the absence of infection, our model tracks the host population size, *N*(*Y*, τ), in year *Y* and within-year time τ, where τ takes values between 0 and 1 corresponding to the start and end of the calendar year, respectively. The population dynamics of the migratory host are described by ordinary differential equations across each stage of the annual cycle [17]. The per-capita mortality rate is assumed to be independent of host density but varies across the annual cycle (μ_j_, where *j* = *b, m, w* denotes breeding, migration, and wintering, respectively). The probability of survival at each stage is given by the following equation:

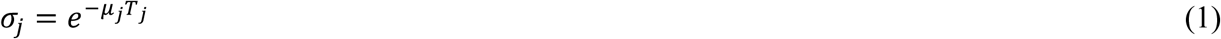

Here, *T*_*j*_ (*j* = *b, m, w*) represents the proportion of the year occupied per stage of the annual cycle. To avoid offspring contributing to reproduction in their hatch year, fecundity is proportional to the number of migrants returning at the start of the breeding season (τ = τ_*b*_), accounting for adult mortality from the onset of the breeding season [50]:

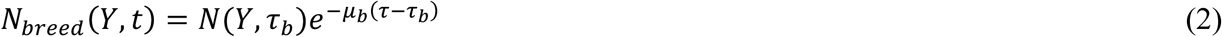

Per-capita reproduction is described by *b*_0_ − *b*_1_*N*, where *b*_0_ and *b*_1_ are the density-independent and density-dependent constants. In the breeding season (i.e., when τ_*b*_ ≤ τ ≤ τ_*b*_ + *T*_*b*_), the population dynamics are as follows:

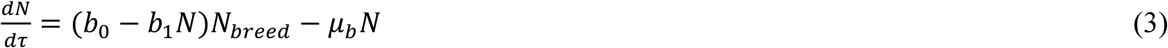

During spring and fall migrations, hosts travel between breeding and wintering grounds each for a duration *T*_*m*_ and experience a per-capita mortality rate μ_*m*_. Hosts spend the remainder of the annual cycle at the wintering grounds (i.e., *T*_*w*_ = 1 − *T*_*b*_ − 2*T*_*m*_) with a per-capita mortality rate μ_*w*_. During migration and winter, population dynamics are given by the following equations:

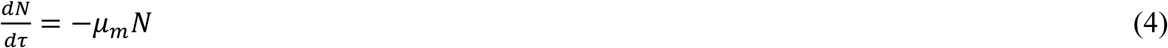

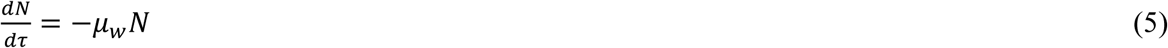

#### Seasonal infection dynamics

To represent the dynamics of relapsing infections, we categorize hosts by their current infection status (i.e., susceptible [*S*], infected [*I*], and latent [*L*]), following the SILI model framework [34,40], where *N* = *S* + *I* + *L*. Transmission from infectious to susceptible hosts in each stage of the annual cycle occurs at the density-dependent rate *β*_*j*_ (see scenarios below), with infected hosts becoming latent at rate ρ (i.e., 1/ρ is the average duration of non-lethal acute infections; Fig. 1). We model stage-specific costs of infection as proportional reductions in host fecundity (*c*_*f*_) and survival (*c*_*j*_), with each ranging from 0 (i.e., no change) to 1 (i.e., infected hosts have a 100% reduction). During the breeding season, we therefore model infection dynamics as follows:

**Figure 1.**
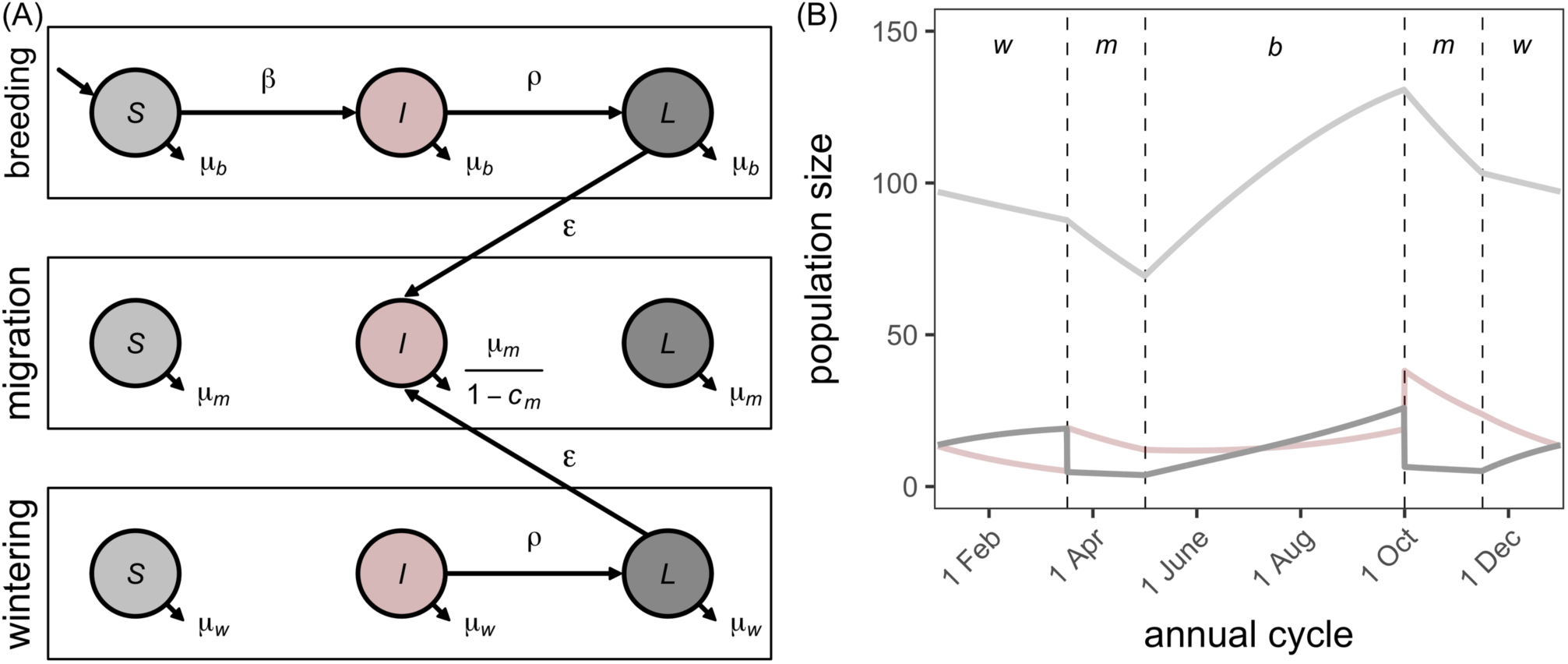
(A) Model schematic of relapsing infections in a migratory host through the annual cycle, for a pathogen transmitted at the host breeding grounds. Filled circles depict the number susceptible (*S*), infectious (*I*) and latently infected (*L*) hosts, and arrows represent gain or losses to each class through demographic or infection processes. Arrows between panels illustrate that a fraction (ε) of latently infected hosts relapse at the start of migration. (B) Example dynamics over the steady state annual cycle. Solid lines depict numbers of susceptible (*S*, light gray), infectious (*I*, pink), and latent (*L*, dark gray) hosts, and vertical dashed lines delineate the spring and fall migratory periods. Infection parameters used are the transmission rate (*β*_*b*_ = 0.05), cost of infection for migrant survival (*c*_*m*_ = 0.5), rate of transition from infectious to latent (ρ = 4, corresponding to a three-month duration of acute infection), and the probability of relapse at the onset of migration (ε = 0.75). Other costs of infection are listed in Table 2 alongside the default values for the demographic parameters.

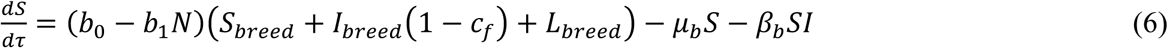

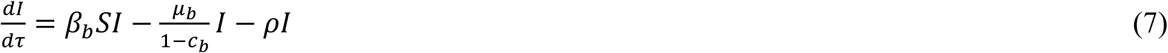

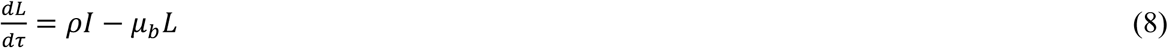

Here, *S*_*breed*_, *I*_*breed*_, and *L*_*breed*_ are the number of returning breeding adults in the susceptible, infected, and latent classes, discounted by their breeding survival (equation 2).

**Table 2.** Model parameters, their definitions, and default values. All rates are given in units of years^-1^. Parameterization for host demography and migration is based on juncos [52–54].

| Parameter | Definition | Value |
| --- | --- | --- |
| Annual cycle |  |  |
| $T_b$ | Proportion of year spent at breeding grounds | 0.42 (~5 months) |
| $T_m$ | Proportion of year spent in each migration | 0.12 (1.5 months) |
| $T_w$ | Proportion of year spent at wintering grounds | 0.34 (~4 months) |
| $\tau_b$ | Start of the breeding season per annual cycle | 0.33 (~early May) |
| Host demography |  |  |
| $b_0$ | Density-independent reproduction rate | 3.3 |
| $b_1$ | Density-dependent reproduction rate | 0.003 |
| $\sigma_b$ | Breeding survival probability | 0.95 |
| $\sigma_m$ | Migration survival probability (one-way) | 0.79 |
| $\sigma_w$ | Winter survival probability | 0.85 (temperate),<br>0.95 (Neotropical) |
| Pathogen traits |  |  |
| $\beta$ | Density-dependent transmission rate | 0.05 (low),<br>0.5 (high) |
| $\rho$ | Infected-to-latent rate | 4 (1 month),<br>12 (3 months) |
| $\varepsilon$ | Fraction of relapsing hosts at the start or end of migration | 0–1 |
| $c_f$ | Cost of infection for fecundity | 0.2 |
| $c_b$ | Cost of infection for breeding survival | 0.2 |
| $c_m$ | Cost of infection for migratory survival | 0.2–0.8 |
| $c_w$ | Cost of infection for winter survival | 0.2 |

Migratory relapse is modeled by ε, representing the fraction of latent hosts that return to the infectious class (Fig. 1). Because latent infections can reactivate shortly after the initiation of migratory restlessness [31], we initially assume relapse occurs instantaneously at the start of spring and fall migrations and is thus driven by physiological preparations for long-distance movement. We subsequently consider an alternative scenario where immunosuppression driven by energetically costly long-distance movement itself results in relapse occurring immediately following migration (i.e., when migrants arrive at their breeding and wintering grounds) [28,51]. At the time of relapse, the number of latent and infected hosts upon each migration are reset:

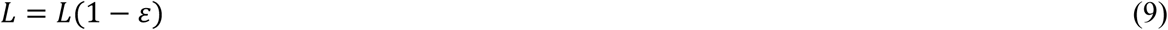

And

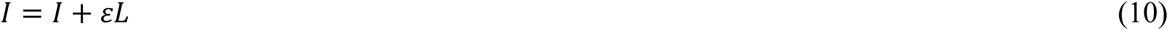

Owing to immunosuppression during migration, we assume acute infections do not subside during migration (i.e., ρ = 0). Infection dynamics during migration are as follows:

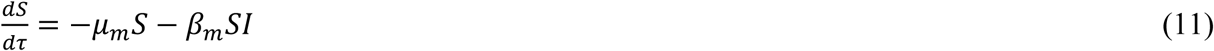

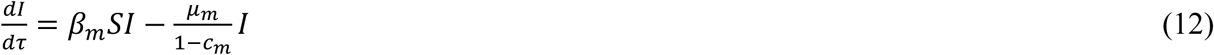

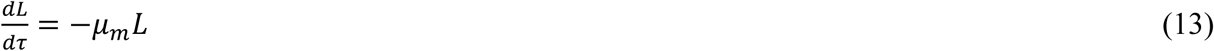

Infection dynamics are modeled in a similar manner at the overwintering grounds (Fig. 1), although infections can here become latent owing to relatively weaker physiological tradeoffs:

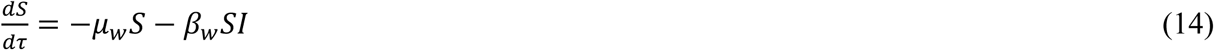

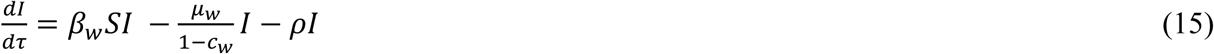

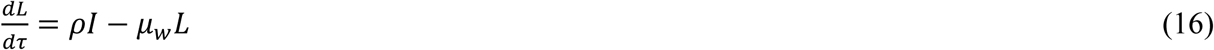

### Parameterization

We parameterized our model using data on the dark-eyed junco (*Junco hyemalis*), a temperate songbird with diverse migratory strategies across North America [52]. Juncos breed at high latitudes and altitudes across Alaska, Canada, and the northern and western United States, migrating south in autumn to a range of overwintering sites. The timing of the junco annual cycle is well-characterized, with migrants beginning to depart their wintering grounds in early March, migrations spanning four to eight weeks, and reproduction starting at the breeding grounds in May (Table 2) [53,54]. Migrants depart their breeding grounds in early October and arrive at the wintering grounds in November. Juncos lay an average of four eggs per clutch across 1–2 broods, resulting in a maximum per-capita fecundity of eight juveniles reared over the five-month breeding season [52]. The survival probability at the breeding grounds is high but is lower during migration. As winter survival probability is higher than that during migration [53], we assume σ_*m*_ < σ_*w*_ < σ_*b*_ to characterize temperate migrants (Table 2). However, we assess sensitivity to this assumption by modeling equal winter and breeding survival probabilities (σ_*w*_ = σ_*b*_), as likely occurs for many Neotropical migrants [55]. Migratory relapse has not been observed in juncos, but this species can be infected with relapsing pathogens (e.g., *B. burgdorferi* and haemosporidians) and displays physiological costs of long-distance migration [19,56,57].

### Infection scenarios and model analysis

Because our main aim was to assess how migratory relapse affects long-term infection prevalence (*I*/*N*), we varied the proportion of hosts that relapse with migration (ε) from 0 (i.e., no reactivation of infection) to 1 (i.e., all latent infections reactivate). In our baseline scenario, we assumed pathogen transmission occurs only at the breeding grounds [17] (i.e., *β*_*b*_ = *β, β*_*m*_ = *β*_*w*_ = 0). This assumption holds for systems where only the breeding season produces enough susceptible hosts to enable transmission or where exposure is driven by breeding behavior [58]. Limiting transmission to the breeding grounds can also coarsely approximate vector-borne diseases given the phenology of many arthropod vectors [32].

To identify pathogen traits that might shape the degree to which migratory relapse affects prevalence, we next systematically varied the duration of acute infection (1/ρ) and transmission rate (*β*) alongside our relapse parameter (ε). Because past work on juncos supports migratory culling in this particular host system [19], and because particularly virulent pathogens could rapidly cull latent hosts that become infected with migratory relapse [1], we also varied the cost of infection for survival during migration (*c*_*m*_). We assumed these costs could be greater than those to fecundity (*c*_*f*_) and survival during breeding (*c*_*b*_) and wintering (*c*_*w*_) stages (Table 2), given that pathogen impacts can be most evident during energetically costly migrations [13,57].

Because host aggregations (e.g., at stopover sites [30]) or particular environmental conditions (e.g., tropical habitats supporting winter activity of vectors [11]) can facilitate some pathogens being transmitted outside of only the breeding season, we explored model behavior across three additional transmission phenologies: (*i*) winter only (*β*_*w*_ = *β, β*_*b*_ = *β*_*m*_ = 0), (*ii*) migration only (*β*_*m*_ = *β, β*_*b*_ = *β*_*w*_ = 0), or (*iii*) year-round (*β*_*b*_ = *β*_*m*_ = *β*_*w*_ = *β*).

Lastly, relapse could occur from not only physiological preparations for migration but also immunosuppression driven by energetically costly long-distance movement itself [28,51], resulting in reactivation occurring closer to the end of their migrations. To account for this alternative timing of relapse, we repeated the above simulations such that the transition from latent to infected occurs when hosts arrive at the breeding and wintering grounds.

Across all parameterizations, we ran our model for at least 25 years and allowed simulations to continue until the maximum mean difference in infection prevalence per timestep with the previous year was below 0.0001, representing a steady state annual cycle (e.g., Fig. 1). Most iterations reached steady state in 40 years. All simulations were conducted in R using the *deSolve* package [59]. For each unique set of model parameters, we recorded the maximum infection prevalence across the steady state annual cycle as a measure of disease risk [50].

## Results

### Migratory relapse and host–pathogen dynamics

In the absence of pathogen relapse (ε = 0) and when transmission occurs only in the breeding season, infection prevalence in the migratory population is relatively low for our baseline parameterization. Prevalence increases early in the breeding season, attains its maximum prior to fall migration, and decreases over the winter through lack of transmission (Fig. 2). In contrast, relapse at the start of migration generally amplifies prevalence across the annual cycle, especially for low-virulence pathogens (i.e., causing a small increase in migratory mortality); at its most extreme (ε = 1), reactivation of all latent hosts upon migration triggers dramatic pulses of infection back into the population (Fig. 2A). Prevalence then declines across the breeding season, mostly as relapsed hosts transition to latency. As more hosts acquire infection, migratory relapse facilitates disease-induced mortality and thereby reduces population size. The largest reductions in population size due to migratory relapse occur for pathogens of intermediate virulence (Fig. 2B). Intermediate infection costs decrease the mean and variance in prevalence throughout the year with migratory relapse across the annual cycle (Fig. 2B). However, when infection poses strong costs for migrant survival and the fraction of reactivation is high, relapse more rapidly converts hosts from latency to actively infected and enhances the effect of migratory culling on infected individuals (i.e., mortality), thereby reducing population-level prevalence (Fig. 2C).

**Figure 2.**
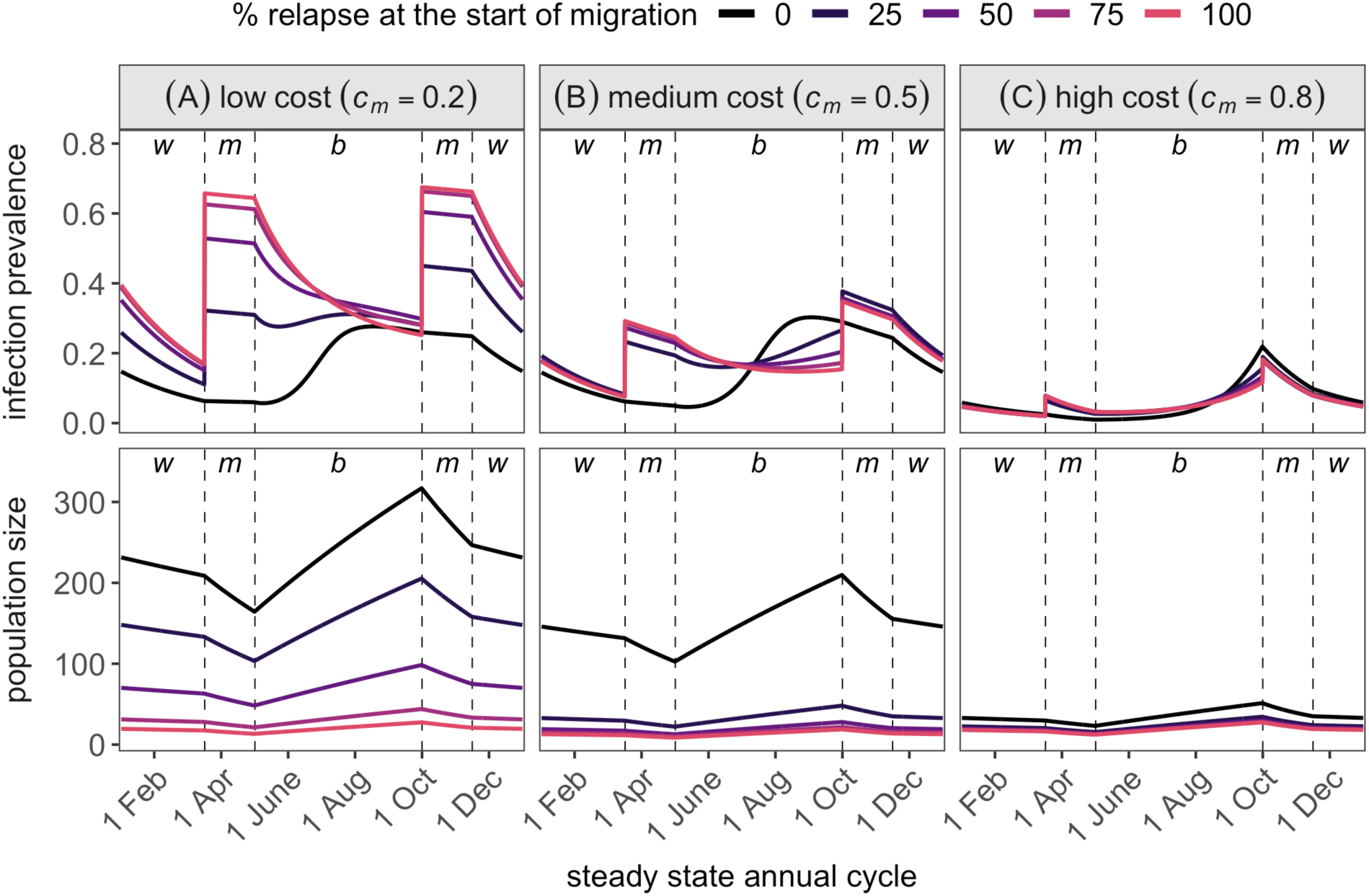
Steady state infection prevalence (top row) and population size (bottom row) across the annual cycle as a function of the fraction of relapse at migration onset (ε, colored lines) and low, medium, and high infection costs for migratory survival (*c*_*m*_, columns). Transmission occurs at the breeding grounds (*β*_*b*_ = 0.5), infection is acute for three months (ρ = 4), and winter survival represents a temperate migrant (σ_*w*_ = 0.85). All other parameter values are listed in Table 2.

### Sensitivity to pathogen traits and host overwinter survival

To understand broader contexts where migratory relapse amplifies or minimizes infection prevalence, we systematically covaried the cost of infection to migratory survival (*c*_*m*_), the fraction of relapsing hosts upon migration (ε), pathogen transmissibility (*β*), and the duration of non-lethal acute infection (1/ρ). We explored patterns for two host migration scenarios: that of a temperate migrant with winter survival lower than breeding survival (σ_*w*_ < σ_*b*_) and that of a Neotropical migrant where winter survival is equivalent to breeding survival (σ_*w*_ = σ_*b*_).

For our baseline (i.e., temperate migrant) scenario, migratory relapse broadly increases the maximum prevalence when costs of infection are low (Fig. 3). For pathogens with low transmissibility (*β* = 0.05) and short acute infections (i.e., one month relative to the six-week migration), migratory relapse allows pathogens to persist that otherwise would be unable to invade the host population (Fig. 3). As expected, pathogens with higher transmissibility and longer acute infections attain greater peak prevalence when costs of infection are low to intermediate. However, migratory relapse can decrease prevalence relative to models assuming no relapse (ε = 0) when costs of infection and transmission rates are both high (Fig. 3).

**Figure 3.**
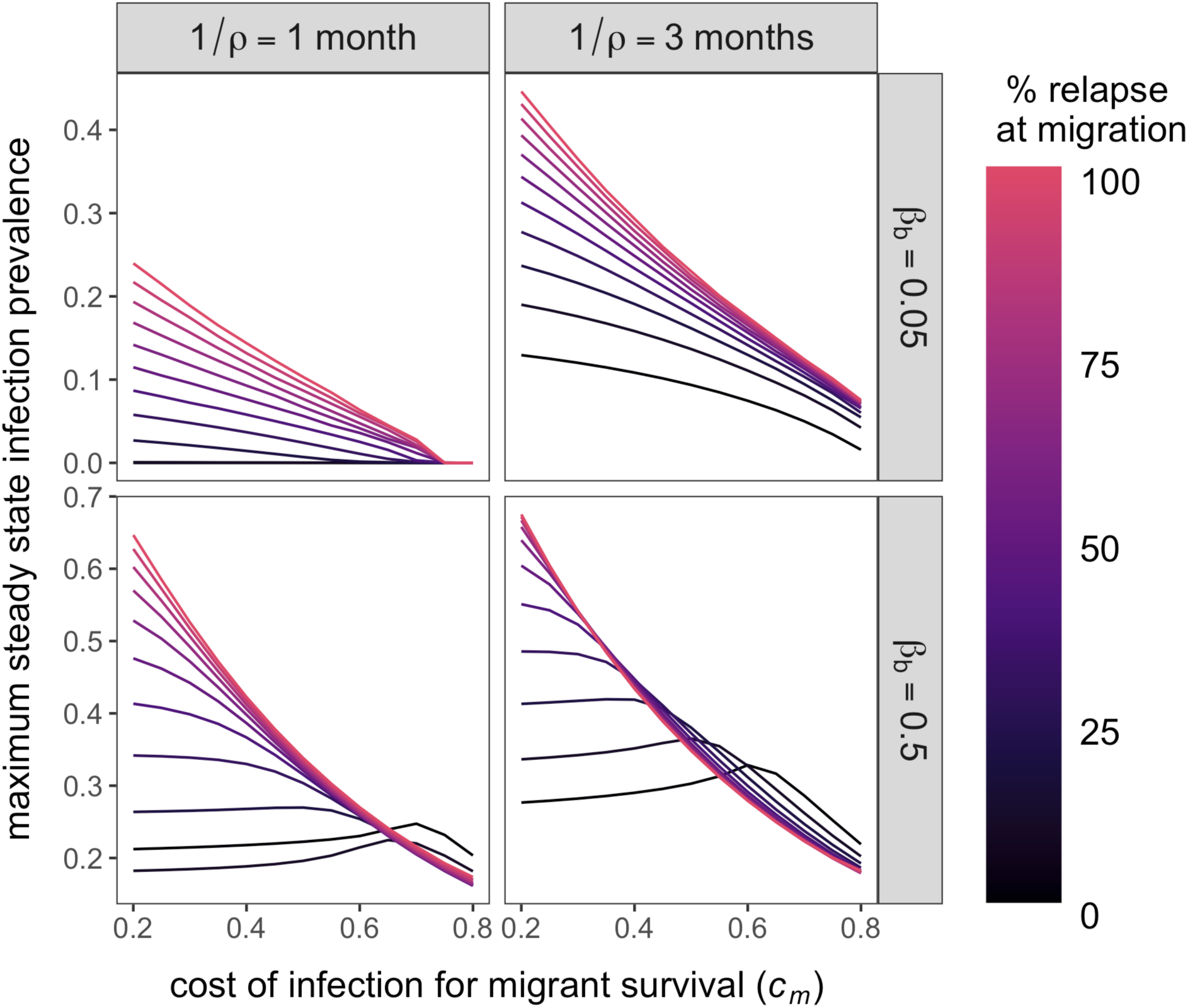
Sensitivity of the relationship between peak infection prevalence and virulence (i.e., cost of infection for migratory survival, *c*_*m*_) to pathogen traits. Transmission occurs only at the breeding grounds with moderate (*β*_*b*_ = 0.05, top) or high (*β*_*b*_ = 0.5, bottom) transmissibility. Lines are colored by the fraction of relapsing hosts at the start of migration (ε), and columns indicate an increasing duration of acute infection (1/ρ) from 1 to 3 months. Winter survival represents a temperate migrant (σ_*w*_ = 0.85), and all other parameter values are listed in Table 2.

These patterns were consistent when winter survival reflected Neotropical migrants (σ_*w*_ = σ_*b*_). Maximum prevalence was overall greater, with the pathogen able to persist with relapse across a broader range of infection costs (Fig. S1). With low costs of infection for migrant survival, migratory relapse produced greater increases in prevalence, particularly with higher transmission rates and longer infectious periods. At higher infection costs, the difference in maximum prevalence between models without relapse (ε = 0) and with complete relapse (ε = 1) was minimized so that migration mostly only dampened infection by virulent pathogens.

### Sensitivity to transmission phenology

We next considered how the timing of pathogen transmission within the migratory host annual cycle affects model outcomes. The amplifying effects of migratory relapse on prevalence are most pronounced when transmission rates are low, irrespective of transmission phenology (Fig. 4). Yet when transmission rates are high, complete migratory relapse (ε = 1) can reduce prevalence relative to a model without relapse (ε = 0) especially when transmission occurs during migration and throughout the annual cycle. For the former, this infection-dampening effect of relapse is particularly pronounced for pathogens with short infectious periods (Fig. S2). These patterns were similar for the Neotropical migrant scenario (i.e. high winter survival), although fewer regions of parameter space reduced infection prevalence with relapse (Fig. S3).

**Figure 4.**
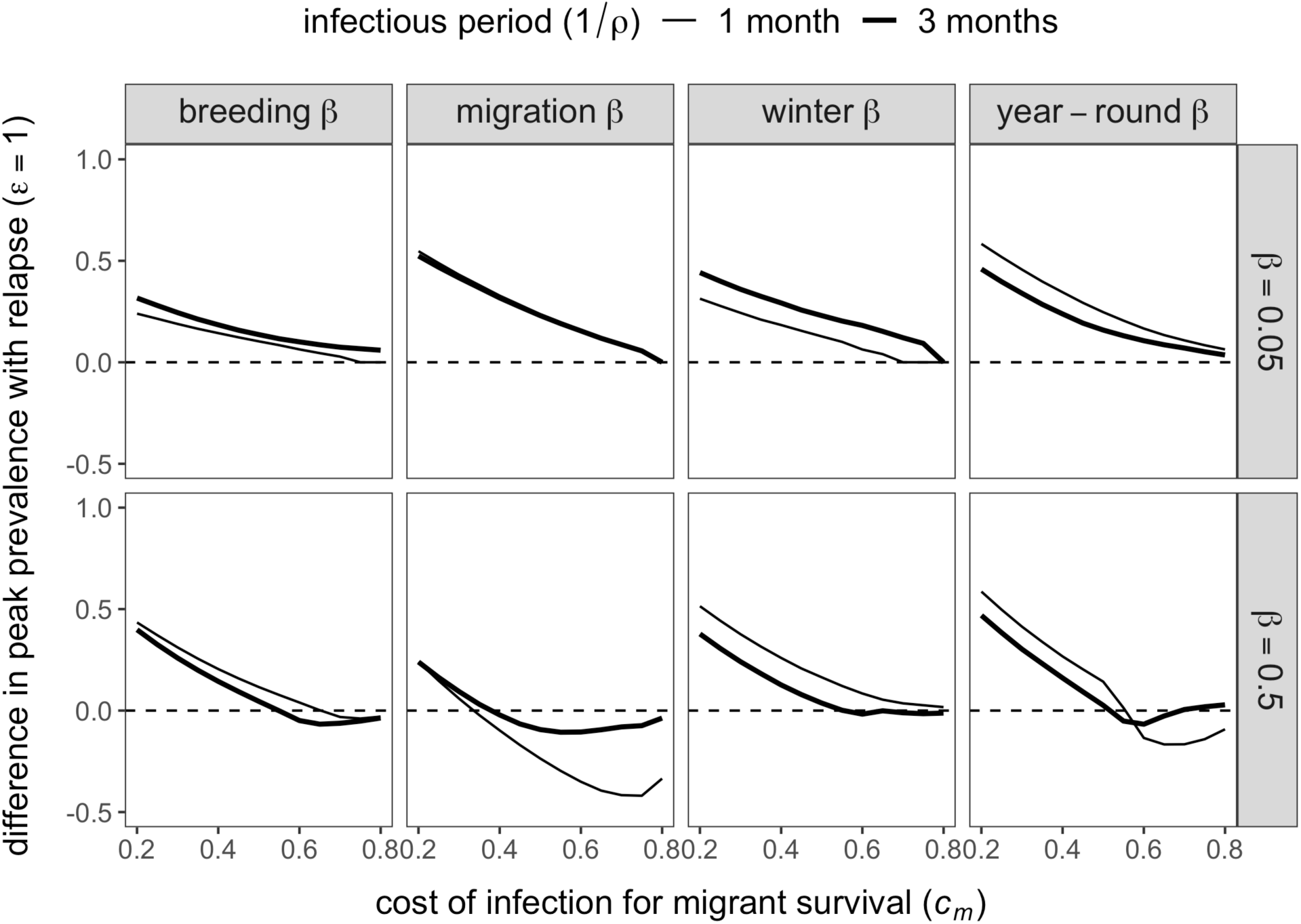
Effects of transmission phenology and pathogen traits on how infection prevalence responds to migratory relapse. We calculate the response of infection prevalence to relapse (vertical axis) as the difference in maximum prevalence between models where all individuals relapse (ε = 1) and no individuals relapse (ε = 0); values above or below the dashed line indicate increases or decreases in peak prevalence with relapse, respectively. Columns represent different transmission phenologies (breeding only, migration only, winter only, or year-round), Rows represent moderate (*β* = 0.05, top) and high (*β* = 0.5, bottom) transmissibility, line width represents short versus long infectious periods (1/ρ), and the horizontal axis represents costs of infection for migrant survival (*c*_*m*_). Winter survival is representative of temperate migrants (σ_*w*_ = 0.85), and all other parameter values are provided in Table 2.

### Sensitivity to the timing of relapse

Lastly, we assessed how relapse at the end of migration, driven by energetically costly long-distance movement itself, affects model outcomes. Because latent hosts undergo relapse after infection costs for survival are most pronounced (i.e., during migration), this later timing of relapse increases maximum prevalence compared to when relapse occurs at the start of migration (Fig. S4). Interestingly, when the cost of infection for migrant survival is high, relapse upon arrival at the breeding grounds amplifies breeding season prevalence enough to then see lower prevalence with relapse during fall migration (Fig. S4). However, broader parameter exploration confirmed that relapse at the end of migration uniformly increases maximum prevalence across our pathogen and host traits (Figs. S5 & S6) and transmission phenologies (Figs. S7 & S8).

## Discussion

Determining the conditions under which migration amplifies or dampens pathogen transmission is important to identify when and where infection risks are greatest in highly mobile species. Migratory species host various pathogens with cycles of latency and reactivation, including several with zoonotic potential [31,40,41], but theory to date on migratory host–pathogen interactions does not account for migration-induced pathogen reactivation and its population- level consequences. Using a mathematical model, we highlight how a novel mechanism, migratory relapse, can increase or decrease infection prevalence throughout the annual cycle, dependent upon pathogen traits and transmission phenology. For pathogens with relatively low virulence, gains to the infectious class through relapse outpace losses of infected individuals through migratory culling to increase prevalence. In contrast, relapse at the start of migration can exacerbate migratory culling to reduce prevalence, primarily for pathogens that are especially virulent, highly transmissible, and spread during the breeding and migratory stages of the annual cycle. By incorporating physiological processes into host–pathogen models (i.e., energetic demands of migration that lead to immunosuppression and relapse [26–28]), our work suggests that within-host mechanisms could account for a wide range of migration–infection patterns.

Past theoretical models of migratory hosts infected by a single pathogen have generally concluded that long-distance movement reduces prevalence by limiting transmission opportunity (i.e., migratory escape), reducing infected host survival (i.e., migratory culling), and increasing host recovery (i.e., migratory recovery) [16–18]. Recent models have also allowed for potential within-host effects of migration during transit through increased recovery (i.e., decreasing prevalence) or increased transmission resulting from immunosuppression [23,60]. However, such models do not examine how these transient effects shape infection patterns across the full annual cycle. Our study illustrates that the effects of migratory relapse can be more important for explaining infection patterns and peak prevalence than direct transmission itself both within and across the annual cycle. In some cases, relapse allowed persistence of pathogens in migrants that would otherwise be purged by insufficient transmission (e.g., due to migratory escape), whereas in other cases relapse eliminated virulent pathogens by increasing migratory culling. Further, in contrast to past models of seasonal infection dynamics, where a single annual peak in prevalence occurs in the transmission season [17], relapse results in a double peak of infection at the beginning or end of each migration. As migrants are often implicated in the transport of zoonotic pathogens such as flaviviruses and influenza viruses [4,29,30,41], this suggests models that fail to incorporate pathogen relapse will not accurately predict when or where peak spillover risk is most likely to occur, with practical implications for pathogen surveillance in migratory species.

For an annual cycle typical of migratory songbirds, and a range of biologically plausible pathogen traits, our model showed that migratory relapse generated dramatic increases in infection prevalence. Although prior theory has not examined relapse induced by physiological changes with migration, past models of relapsing pathogens have demonstrated that reactivation can facilitate pathogen persistence. In the absence of immigration, recurrent cycles of acute infection and latency were necessary to explain henipavirus dynamics in straw-colored fruit bats [40], and more frequent relapse optimized pathogen invasion potential for human malaria (*Plasmodium vivax*) [45]. Our model suggests migratory relapse can generate pronounced biannual peaks in prevalence that maintain more infected hosts throughout the annual cycle than direct transmission alone. This biannual pulse differs in its seasonal timing and magnitude from those generated by seasonal birth pulses or relapse induced by immune tradeoffs with reproduction [37,39,61,62], especially in systems where offspring benefit from prolonged maternal immunity [63]. Because this pattern was most evident when infection had low costs for migrant survival, our predictions may be especially applicable to migratory reservoir hosts that themselves do not experience high morbidity or mortality (e.g., henipaviruses in flying foxes and *Borrelia burgdorferi* in songbirds [31,40]). Given the high prevalence of actively infectious hosts during migration, stopover sites and periods of migrants returning to breeding or wintering grounds could be prioritized as important targets for zoonotic pathogen surveillance [64].

Our model also identified contexts in which migratory relapse can decrease prevalence, or even cause pathogen extinction, by removing more actively infectious hosts with migratory culling. Reductions in prevalence were most likely to be seen in highly transmissible pathogens, those with long infectious periods, and when transmission occurred at the breeding grounds or during migration. Systems for which such conditions are met include rhabdoviruses in salmonid fish as well as haemosporidian parasites and flaviviruses in some songbirds [7,19,51,65]. High relapse frequency and high mortality of infected migrants might be more common for species undertaking strenuous migrations, such as those involving prolonged periods of powered flight. This supports prior predictions that migratory culling is more likely to operate in long-distance migrations with fewer stopovers than shorter migrations or where migratory species move and forage in a stable environmental window (e.g., ungulates “surfing the green wave”) [17,66].

Given the sensitivity of our model to the degree of relapse and the costs of infection for migrant survival, estimation of these parameters will be important for applying this framework to empirical systems. Detecting pathogen relapse requires sampling hosts across their annual cycle [34], which can be particularly logistically challenging for highly mobile species [67]. However, advancements in tracking technologies (e.g., lightweight geolocators, stable isotopes) are improving inference into migratory networks, which could facilitate temporal sampling across breeding, wintering, and stopover habitats [68]. Additionally, multiple diagnostic efforts are necessary to differentiate compartments within the SILI framework. For example, serology, PCR, and microscopy can distinguish uninfected (i.e., susceptible), acutely infected, and latent hosts for haemosporidian parasites [69], and pathogen genotyping could disentangle whether infections are more likely to indicate reactivation rather than novel transmission events [34]. Integrating longitudinal sampling and these diagnostics with mark–recapture could help to estimate infection- and stage-specific survival probabilities and thus infer pathogen costs [70], which would be especially important for more realistic models (e.g., explicit age structure).

To examine conditions under which relapse during migration increases or decreases infection risks, we focused our model on a simple system with a single interaction between a migratory host and its pathogen. For tractability, we ignored alternative ecological interactions such as multiple host species (and thus a more generalist pathogen) or explicit arthropod vectors. However, high prevalence in relapsing migrants suggests migrants could introduce pulses of infection into seasonally sympatric co-occurring species. This could be particularly important for pathogen maintenance in partially migratory species such as juncos, where long-distance migrants form mixed flocks with residents at shared wintering grounds [52]. For vector-borne pathogens with cycles of latency and reactivation, such as *Borrelia burgdorferi* and some flaviviruses [31,71], migrants undergoing relapse could also have disproportionate influence on infecting vectors upon their arrival at the breeding grounds. Further, as some infections may ultimately reduce migratory propensity itself [72], relapse may drive transitions to residency. Future extensions to our model could assess how phenological overlap between migrants and vectors shapes disease outcomes as well as feedbacks between relapse and migration [50,73].

Although our model incorporates within-host processes into the population-level dynamics of infection among migratory animals, we also only consider a simplified system in which long-distance movement impairs immune defense through physiological tradeoffs. Such an assumption is supported by comparative and experimental studies of migration and immunity [26–28]; however, despite clear energetic costs of migration, some hosts can maintain equivalent immune function during migration or even enhance specific immune responses as an evolved mechanism to increase survival [74,75]. Explicitly modeling how particular immune axes are up- or downregulated prior to and during migration could better inform how host susceptibility and relapse vary across the annual cycle and their consequences for infectious disease dynamics.

Lastly, because many species are shifting the timing and extent of their migration in response to climate and anthropogenic change [9,76], associated changes to host physiology and survival could alter how migratory relapse affects infection patterns. Deterioration of resources at breeding or wintering sites could increase the proportion of relapsing hosts, increasing prevalence of low-virulence pathogens or preventing persistence of high-virulence pathogens if migratory mortality is also elevated. Additionally, many migratory species are increasingly overwintering in anthropogenic habitats. For example, Anna’s hummingbirds and European blackcaps both spend more of their annual cycle in urbanized regions owing to abundant and predictable anthropogenic food resources [77,78]. Such resources could dampen or amplify the likelihood of pathogen relapse at spring departure depending on their nutritional quality or effects on host density and crowding [79]. Other stressors associated with urban habitats, such as artificial light at night, could further amplify the likelihood of relapse [80,81]. These consequences of urban habituation could be especially relevant for human health in the context of wildlife reservoirs of relapsing zoonoses, such as flying foxes and henipaviruses [82].

Many pathogens of concern to human, domestic animal, and wildlife health exhibit cycles of latency and reactivation. Our model provides a generalizable and mechanistic framework for understanding the dynamics of such infections in migratory hosts. We demonstrate theoretical support for migratory relapse increasing infectious disease risks but also emphasize the context dependence of these patterns on host and pathogen traits. In particular, we illustrate that this within-host process can generate both increases and decreases in infection prevalence from migration, providing a mechanism for explaining divergent associations between migration and disease in empirical systems. Empirical estimates of relapse and survival across the annual cycle will be important for linking such frameworks with empirical systems, which will be critical for prioritizing pathogen surveillance in migratory species and predicting how changes in climate and land use alter migration, reactivation of latent infections, and pathogen spillover risks.

## Data accessibility

R code for reproducing the main analyses is available in the Dryad Digital Repository: https://datadryad.org/stash/share/eT4DN6oJ0BlGmv5fqjg16cm3HThgXmvgllWemPpInYY.

## Author contributions

DJB and RJH designed the model, DJB and EDK parameterized the model, and DJB performed the analyses and wrote the first draft of the manuscript. All authors contributed to revisions.

## Acknowledgements

We thank members of the Ketterson lab and two reviewers for constructive feedback.

## Funding

DJB was supported by an appointment to the Intelligence Community Postdoctoral Research Fellowship Program, administered by Oak Ridge Institute for Science and Education through an interagency agreement between the U.S. Department of Energy and the Office of the Director of National Intelligence. EDK was supported by the Environmental Resilience Institute, funded by Indiana University’s Prepared for Environmental Change Grand Challenge Initiative. RJH was supported by the National Science Foundation (DEB-1754392 and DEB-1911925).

## Competing interests

We declare no competing interests.

## Notes

### Competing Interest Statement

The authors have declared no competing interest.

