## Supplemental Material for "Reactivation of latent infections with migration shapes population-level disease dynamics"

Daniel J. Becker, Ellen D. Ketterson, Richard J. Hall

Figure S1. Sensitivity of peak infection prevalence to pathogen traits. We varied the fraction of relapsing hosts at the start of migration ($\varepsilon$) and the costs of infection for migrant survival ($c_m$) for select values of transmissibility ($\beta$) and the duration of acute infection ($1/\rho$, shown as months). Maximum prevalence is displayed for each parameterization under Neotropical migrant assumptions in which winter survival is equivalent to breeding survival ($\sigma_w = \sigma_b = 0.95$).

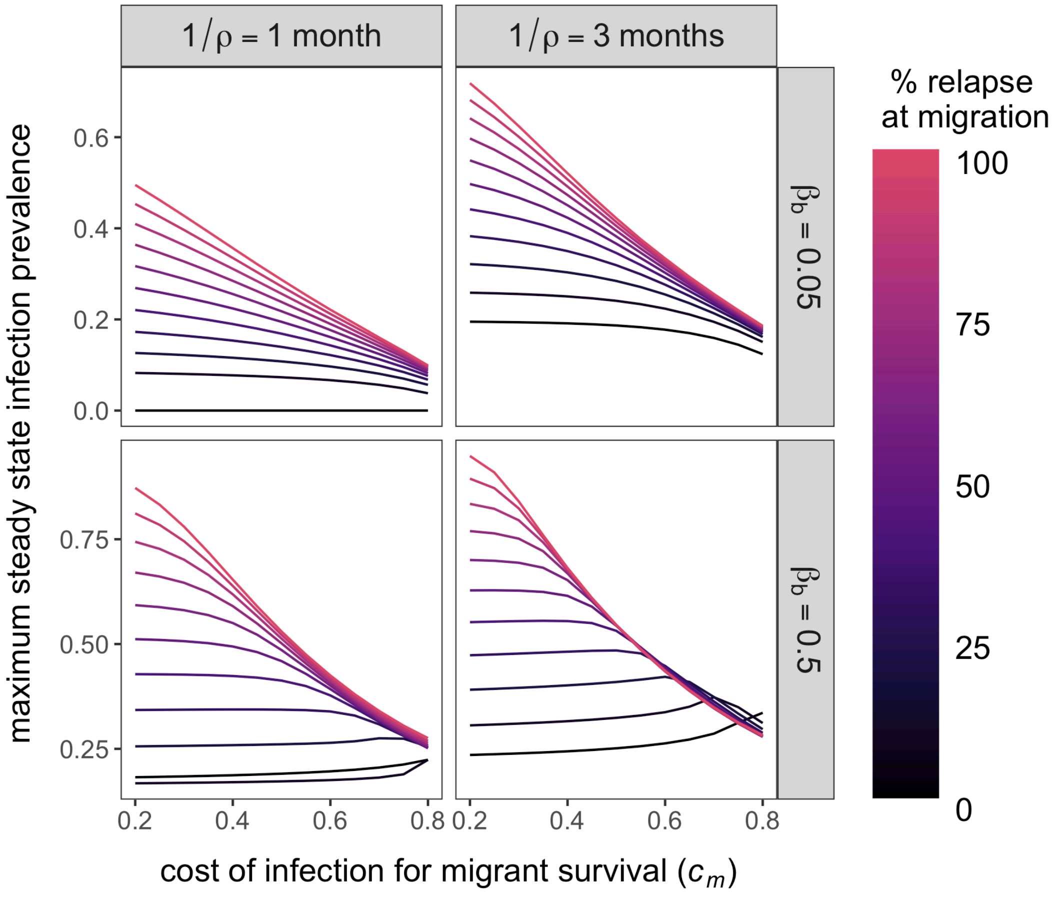

Figure S2. Sensitivity of peak infection prevalence to pathogen traits when transmission only occurs during migration. As in the main text, we varied the fraction of relapsing hosts at the start of migration ($\varepsilon$) and the costs of infection for migrant survival ($c_m$) for select values of transmissibility ($\beta$) and the duration of acute infection ($1/\rho$, shown as months). Maximum prevalence is shown for under temperate migrant survival assumptions ($\sigma_w = 0.85$).

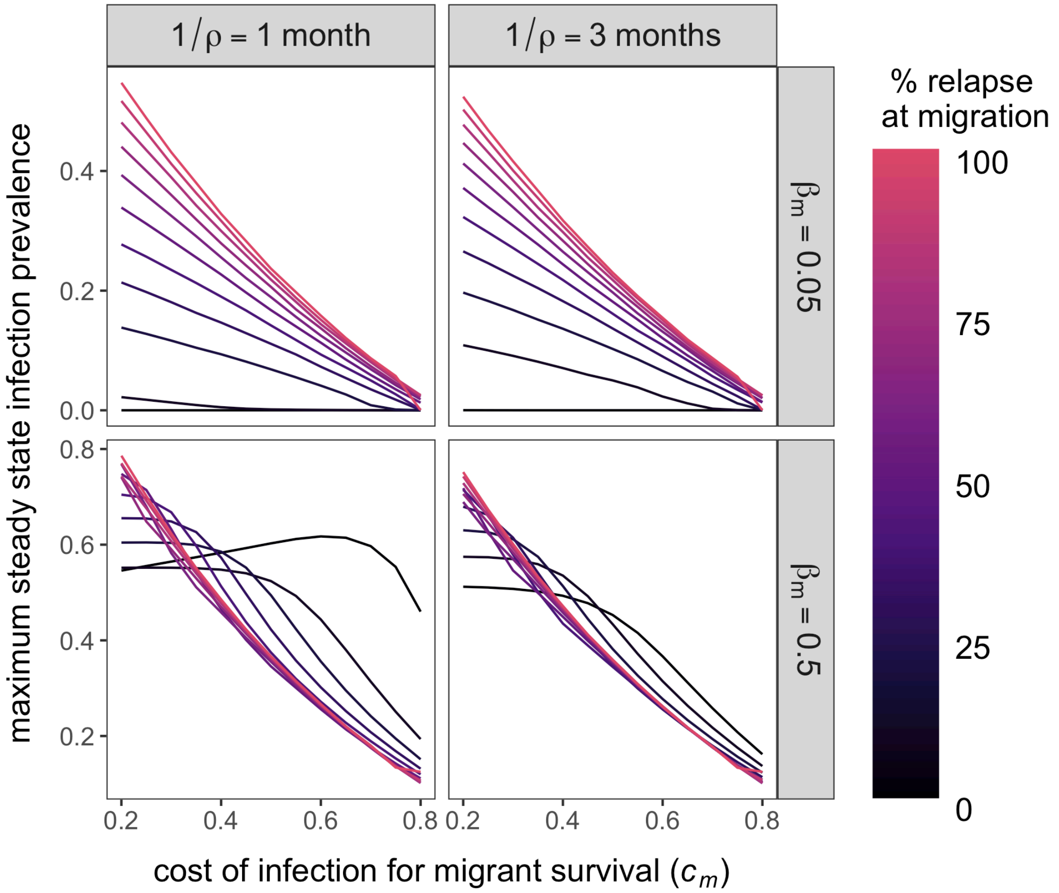

Figure S3. Effects of transmission phenology and pathogen traits on the response to prevalence to migratory relapse. The difference in maximum prevalence between a null model ($\varepsilon = 0$) and maximum relapse ($\varepsilon = 1$) is shown across the range of infection costs for migrant survival ($c_m$), infectious periods ($1/\rho$, shown as months), and transmission rates ($\beta$). We consider our baseline model (transmission in the breeding season only) alongside transmission during migration only, at the wintering ground only, and across all stages of the annual cycle. The dashed line indicates no difference in maximum prevalence, with values above and below corresponding to migratory relapse increasing or decreasing infection. Results are shown under Neotropical migrant assumptions in which winter survival is equivalent to breeding survival ($\sigma_w = \sigma_b = 0.95$).

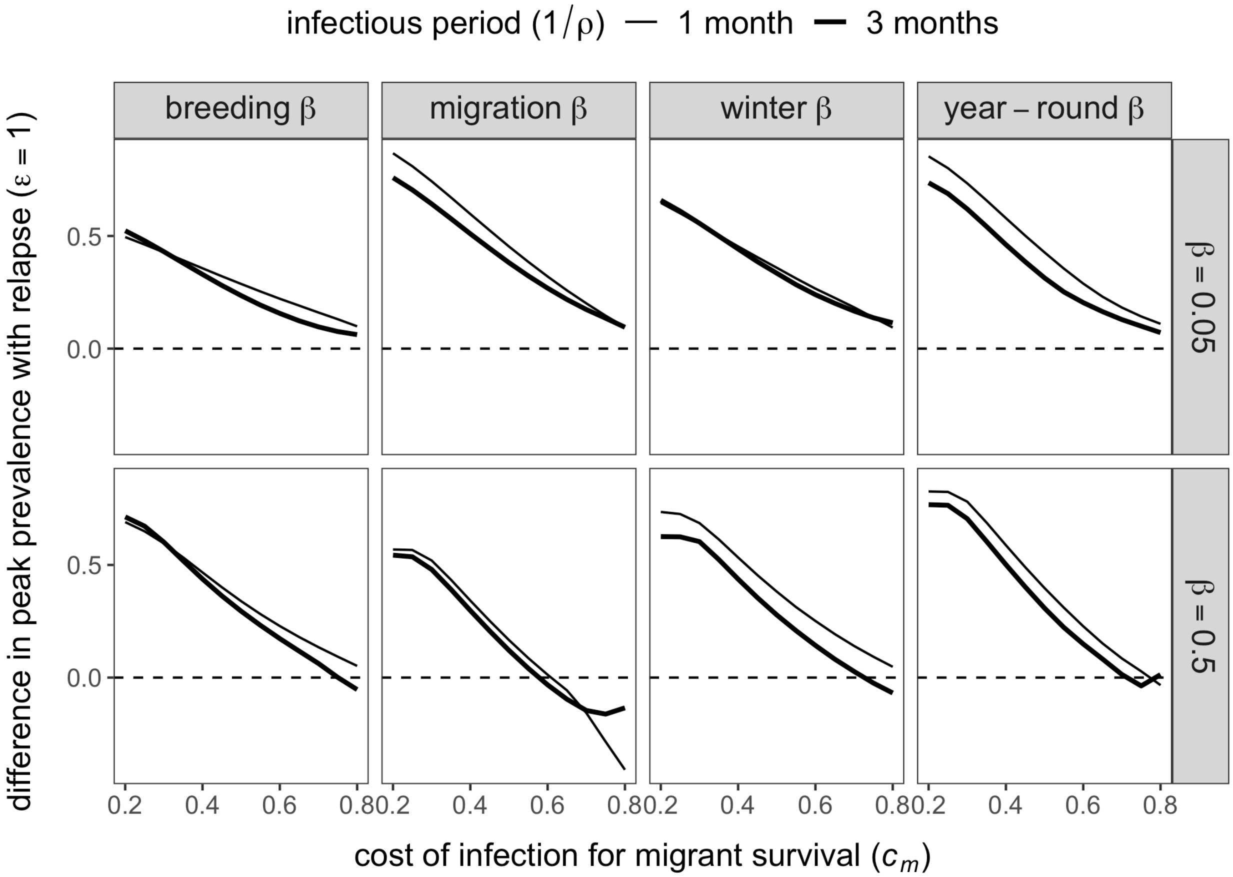

Figure S4. Steady state infection prevalence across the annual cycle as a function of the fraction of hosts that relapse ($\varepsilon$, colored lines) at the start (top row) and end (bottom row) of migration, for low, medium, and high infection costs for migrant survival ($c_m$, columns). Transmission occurs at the breeding grounds ($\beta_b = 0.5$), infection is acute for three months ($\rho = 4$), and winter survival represents a temperate migrant ($\sigma_w = 0.85$). Other parameters are in Table 2.

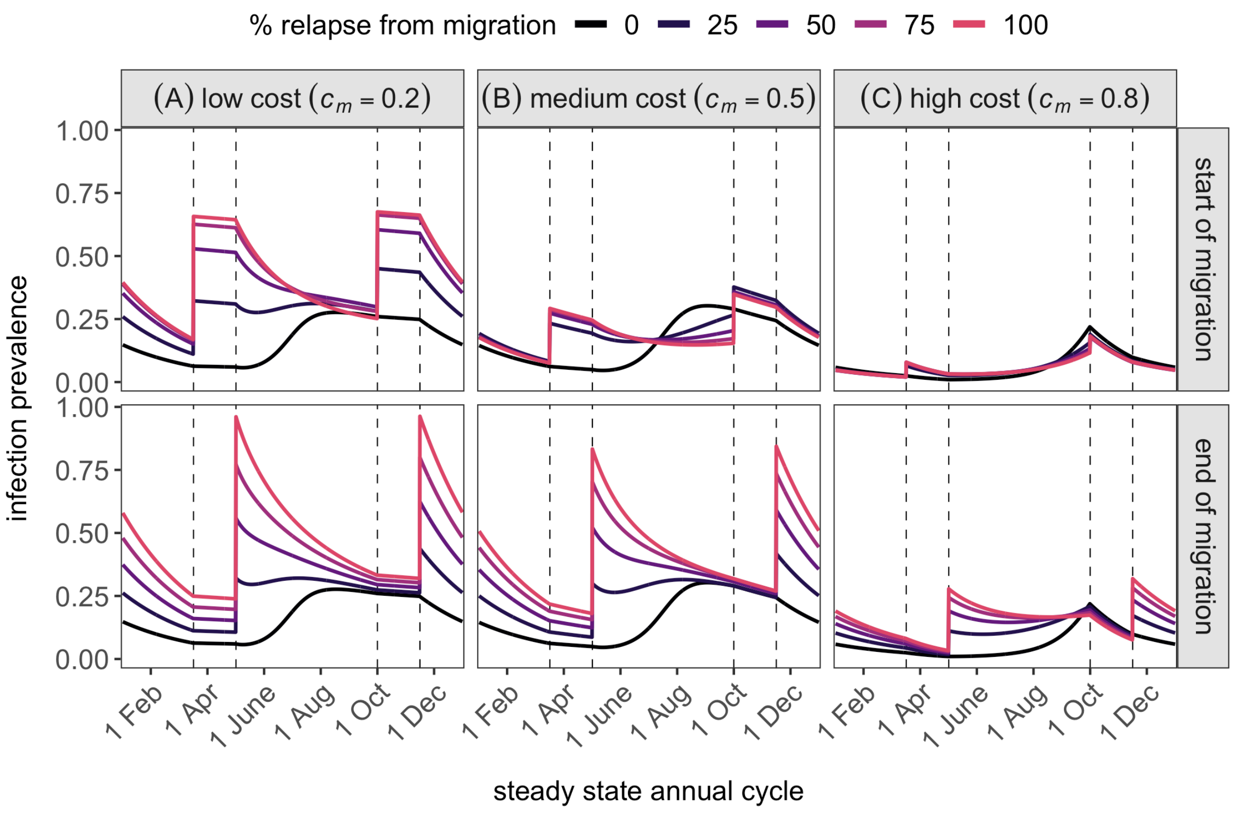

Figure S5. Sensitivity of maximum infection prevalence to pathogen traits when relapse occurs at the end of migration (i.e., upon migrant arrival). We varied the fraction of relapsing hosts ($\varepsilon$) and the costs of infection for survival during migration ($c_m$) for select values of transmissibility ($\beta$) and the duration of acute infection ($1/\rho$, shown as months). Winter survival represents a temperate migrant ($\sigma_w = 0.85$), and all other parameter values are listed in Table 2.

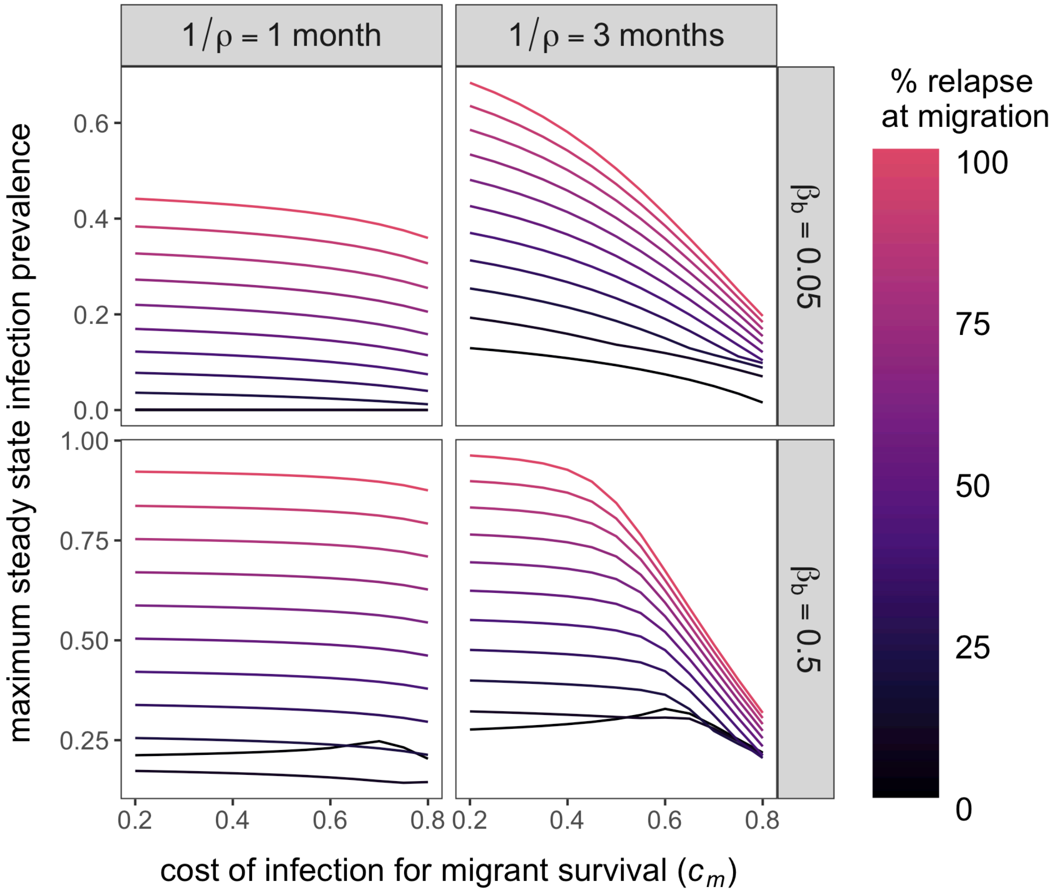

Figure S6. Sensitivity of maximum infection prevalence to pathogen traits when relapse occurs at the end of migration (i.e., upon migrant arrival). We varied the fraction of relapsing hosts ($\varepsilon$) and the costs of infection for survival during migration ($c_m$) for select values of transmissibility ($\beta$) and the duration of acute infection ($1/\rho$, shown as months). Winter survival represents a Neotropical migrant ($\sigma_w = \sigma_b = 0.95$), and all other parameter values are listed in Table 2.

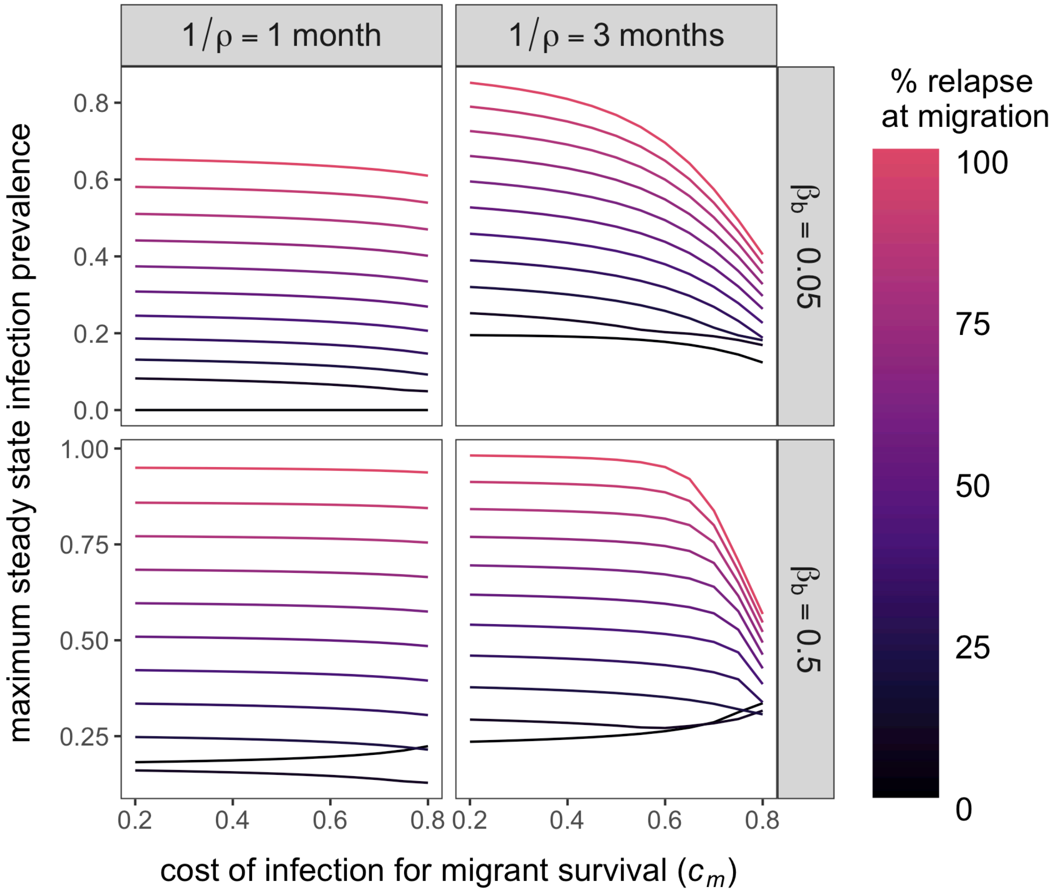

Figure S7. Effects of transmission phenology and pathogen traits on the response to prevalence to migratory relapse. The difference in maximum prevalence between a null model ($\varepsilon = 0$) and maximum relapse ($\varepsilon = 1$) is shown across the range of infection costs for migrant survival ($c_m$), infectious periods ($1/\rho$, shown as months), and transmission rates ($\beta$). We consider our baseline model (transmission in the breeding season only) alongside transmission during migration only, at the wintering ground only, and across all stages of the annual cycle. The dashed line indicates no difference in maximum prevalence, with values above and below corresponding to migratory relapse increasing or decreasing infection. Results are shown under temperate migrant assumptions in which winter survival is lower than breeding survival ($\sigma_w = 0.85$).

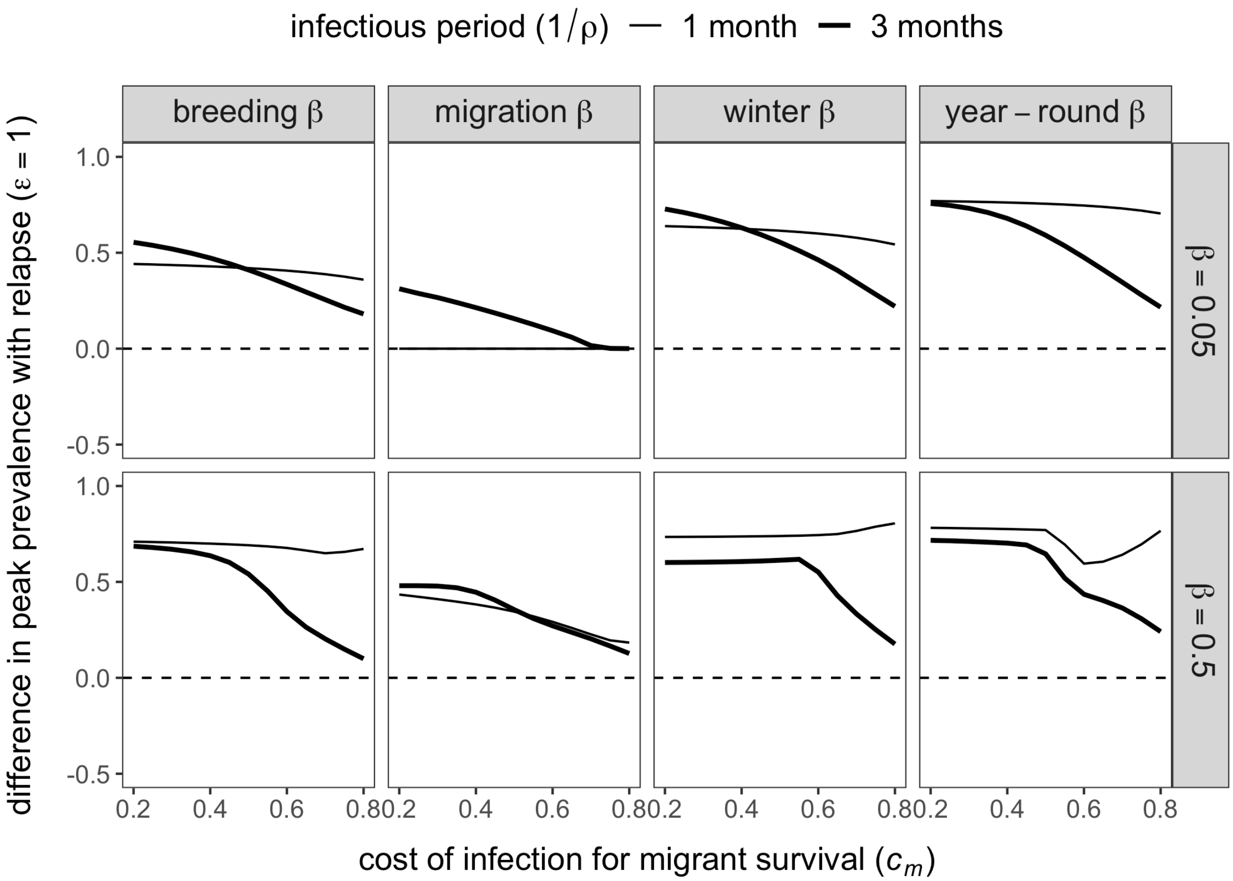

Figure S8. Effects of transmission phenology and pathogen traits on the response to prevalence to migratory relapse. The difference in maximum prevalence between a null model ($\varepsilon = 0$) and maximum relapse ($\varepsilon = 1$) is shown across the range of infection costs for migrant survival ($c_m$), infectious periods ($1/\rho$, shown as months), and transmission rates ($\beta$). We consider our baseline model (transmission in the breeding season only) alongside transmission during migration only, at the wintering ground only, and across all stages of the annual cycle. The dashed line indicates no difference in maximum prevalence, with values above and below corresponding to migratory relapse increasing or decreasing infection. Results are shown under Neotropical migrant assumptions in which winter survival is equivalent to breeding survival ($\sigma_w = \sigma_b = 0.95$).

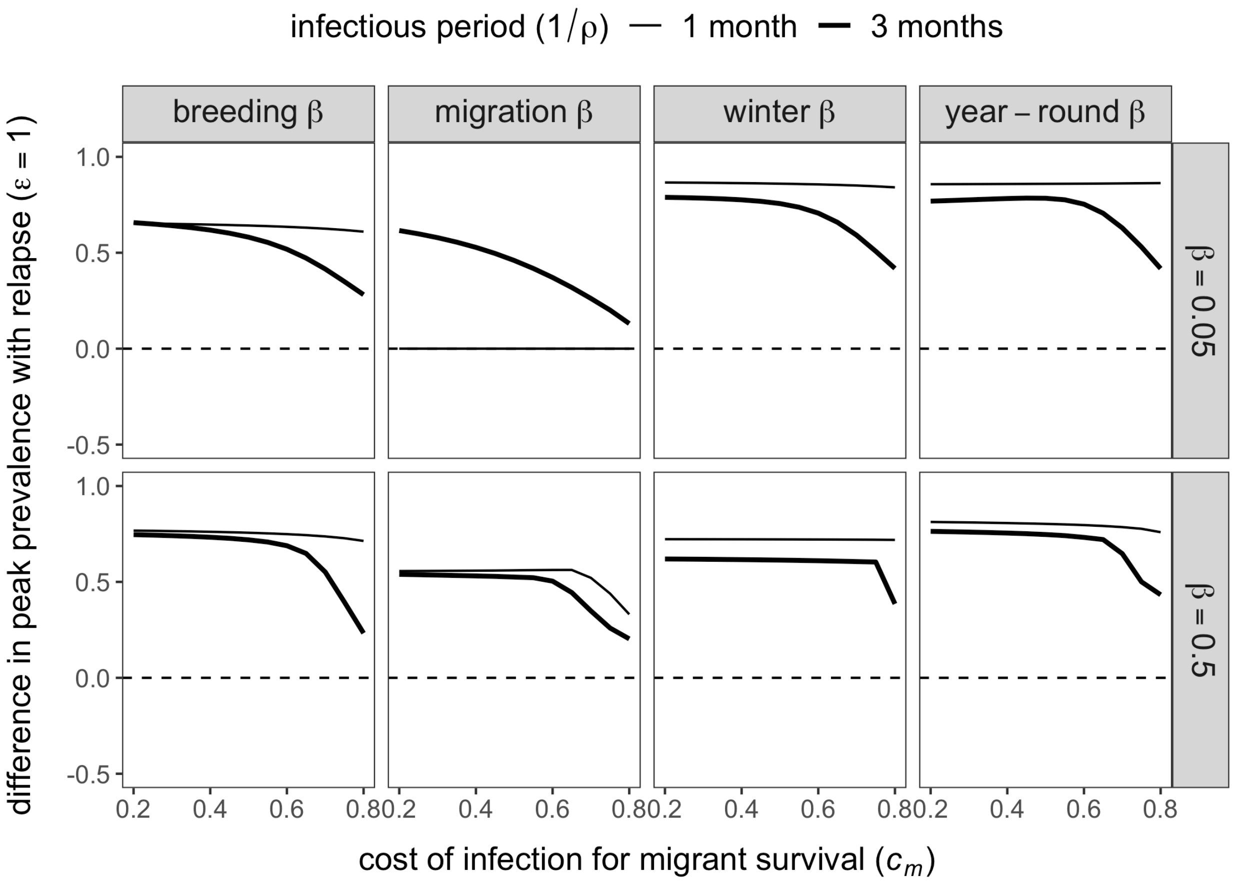
